# Exaggerated postnatal surge of orexin and the effects of elimination of excess orexin on blood pressure in spontaneously hypertensive rats in postnatal development

**DOI:** 10.1101/2020.06.17.158279

**Authors:** Savannah Barnett, Ruhong Dong, Logan Briggs, Alexander Moushey, Aihua Li

## Abstract

It has been established that an overactive orexin (OX) system is associated with neurogenic hypertension in spontaneously hypertensive rats (SHRs). However, the chronology and mechanism of such association between orexin system and hypertension is unclear. We hypothesized that an aberrant surge of OX neurons in SHRs precedes the aberrant increase of arterial blood pressure (ABP) during postnatal development, which was primarily contributed by the exaggerated postnatal OX neurogenesis. We found that (1) SHRs experienced a greater surge in the number of orexin neurons than normotensive Wistar-Kyoto (WKY) rats before P16, which led to significantly more OX neurons than age-matched controls by P15-16 (3680±219 vs 2407±182, respectively, P=0.002). (2) Exaggerated OX neurogenesis, marked by bromodeoxyuridine (BrdU), was the primary contributor to excessive OX neurons in SHRs during development. (3) In contrast, SHRs and normotensive control rats have similar mean arterial blood pressure (ABP) at P15, and a significantly higher ABP in SHR than WKY emerges at P20 (74.8 ± 2.5 vs 66.9 ± 4.4 mmHg in wakefulness, respectively, P<0.05), a few days following the surge of OX activity. (4) Selectively eliminating excess (∼30%) orexin neurons, via a targeted neurotoxin, in SHRs between P30 and P40 results in a significantly lowered ABP compared to non-lesioned SHRs at P40. We suggest that the postnatal surge of OX neurons, primarily attributed to the exaggerated postnatal OX neurogenesis, may be necessary for the development of higher ABP in SHRs, and modulation of the overactive OX system may have a preventative effect during the pre-hypertensive period.

**New Findings:** *What is the central question of this study?:* Excess orexin neurons have been associated with hypertension in spontaneously hypertensive rats, however, the association and mechanism between developing excess orexin neurons and high blood pressure are unknown.

*What is the main finding and its importance?:* Using spontaneously hypertensive rats in anatomical and physiological studies, we provided evidence showing that the excess OX neurons, primarily via exaggerated OX neurogenesis, may be necessary in developing a higher ABP in SHRs during development, and modulation of the overactive orexin system may be beneficial in treating hypertension.

## Introduction

Participation of hypothalamic orexin neurons in cardiorespiratory regulation has been well established over the last two decades (Samson *et al.*, 1999; Shirasaka *et al.*, 1999; Shahid *et al.*, 2011; Li & Nattie, 2014). Orexin neurons project to and excite many cardiorespiratory control nuclei in the central nervous system, including the paraventricular nucleus (PVN), nucleus tractus solitarius (NTS), retrotrapezoidal nucleus (RTN), medullary raphe, rostro-ventrolateral medulla (RVLM), and sympathetic preganglionic neurons of the intermediolateral column (Peyron *et al.*, 1998; Trivedi *et al.*, 1998; Van Den Pol, 1999; Antunes *et al.*, 2001; Marcus *et al.*, 2001; Shirasaka *et al.*, 2001; Yang & Ferguson, 2003; van den Top *et al.*, 2003; Huang *et al.*, 2010; Shahid *et al.*, 2012). Central administration of orexin (OX) leads to significant and sustained increases in arterial blood pressure (ABP), heart rate, catecholamine release, and renal sympathetic nerve activity (RSNA) in conscious and anesthetized rats, an effect that is prevented by pre-treatment with OXR antagonists (Samson *et al.*, 1999; Shirasaka *et al.*, 1999; Huang 2010; Shahid *et al.*, 2011; Li & Nattie, 2014).

A link between an overactive orexin system and the pathophysiology of neurogenic hypertension has been recently established in spontaneously hypertensive rats (SHRs) and mice (Lee *et al.*, 2013, 2015; Li *et al.*, 2013; Clifford *et al.*, 2015; Jackson *et al.*, 2016). On average, adult spontaneously hypertensive mice and rats have about 30% more orexin-producing neurons than their normotensive background controls (Lee *et al.*, 2015; Li *et al.*, 2016; Jackson *et al.*, 2016). Modulating the overactive orexin system via blocking both orexin receptors with a dual orexin receptor antagonist can 1) significantly lower ABP (by 20-30 mmHg), sympathetic nerve activity (SNA), and catecholamine release in conscious adult SHRs (Li *et al.*, 2013) and normalize ABP in younger SHRs (postnatal day (P)30-58), and 2) normalize the exaggerated ventilatory hypercapnic chemoreflex at both ages (Li *et al.*, 2016). These findings suggest that excessive orexin-producing neurons may play important roles in the pathophysiology of neurogenic hypertension in SHRs including the regulation of both the ABP and hypercapnic ventilatory chemoreflex. At present, the chronology of the association between an overactive orexin system and development of hypertension, the sources of excess orexin neurons, and the mechanism of this relationship remain unclear.

The orexin system encounters pre- and post-natal developmental changes and matures postnatally in mammals (Yamamoto *et al.*, 2000; Steininger *et al.*, 2004; Amiot *et al.*, 2005; Sawai *et al.*, 2010; Stoyanova *et al.*, 2010; Iwasa *et al.*, 2015). Using *in situ* hybridization, Yamamoto *et al.* showed that *prepro-orexin* mRNA in the rat hypothalamus gradually increased during early postnatal development and peaked by 3 weeks of age (Yamamoto *et al.*, 2000). Further, Sawai *et al.* showed a clear increase in the number of OXA- and OXB-peptide-expressing neurons from 2 weeks to 9 weeks of age (Sawai *et al.*, 2010). It is not clear whether the postnatal increase in orexin was due to postnatal cell maturation or new cell proliferation, e.g. neurogenesis. Additionally, these studies were completed in normotensive animals and did not include comprehensive cell counts at various ages. At present, little is known about whether the increased number of OX neurons in SHRs is developed prenatally or postnatally, how orexin neuron development and blood pressure development are chronologically correlated postnatally, and whether or not modulation of the OX system can change the course of developing higher ABP during development.

In this study, using SHR and their normotensive background control Wistar–Kyoto (WKY) rat pups, we aimed to address the following questions: 1) At which developmental age the ABP and number of orexin neurons in SHRs deviate from age-matched normotensive WKY rat pups; 2) What is the primary contributing factor to the excess proliferation of orexin-producing neurons during postnatal development in SHRs e.g. neurogenesis or cell maturation; 3) What is the chronological relationship between excess orexin neurons and the development of hypertension in SHRs and WKYs; and 4) Does elimination of excess orexin neurons have an impact on the development of a higher ABP and exaggerated hypercapnic chemoreflex during development. We hypothesized that exaggerated postnatal orexin neuron neurogenesis leads to a postnatal surge of orexin neurons in SHRs, which facilitate an aberrant increase in ABP in SHRs, and that elimination of these excess orexin neurons can prevent SHRs from developing a higher ABP and hypercapnic chemoreflex during a developing age.

## Materials and Methods

### Ethical approval

All animal experimental and surgical protocols were within the guidelines of the National Institutes of Health for animal use and care and were approved by the Institutional Animal Care and Use Committee at the Geisel School of Medicine at Dartmouth. All animals were reared and transported in accordance with regulations of these governing bodies.

### General methods

SHRs and normotensive WKY rats were used for the experiments in this study. All rats were housed in a temperature- and light-controlled environment set on a 12h-12h light-dark cycle (lights on at 00.00 h; lights off at 12.00 h). Food and water were provided *ad libitum*. The general methods are those in common use in our laboratory (Li *et al.*, 2008, 2013, 2016). A total of 75 SHRs and 44 WKY rats between the ages of P7 and P40 were used in four sets of anatomical and physiological experiments, and males and females were assigned into each group randomly. At the conclusion of the experiments, the rats were euthanized with an overdose of sodium pentobarbital (>75 mg kg^-1^, i.p.; Euthasol; Virbac Inc., Fort Worth, TX, USA).

### Surgeries and procedures

#### Blood pressure probe and EEG/ EMG

All animals used to study ABP were surgically implanted with a telemetric blood pressure probe. Rats were anesthetized with isoflurane (Piramal Enterprises ltd.) or a ketamine (Putney, Inc., Portland, ME, USA) and xylazine (Lloyd Labs, Walnut, CA, USA) cocktail (90/15 mg kg^-1^, I.M.). Ketoprofen (3 mg/kg, subcutaneously) was used as an analgesic after surgery. All rats were implanted with an HD-X11 or PA-C10 telemetric probe (DSI, St. Paul, MN, USA), which allows for uninterrupted and reliable recording of ABP during the experiment as similarly described in our previous publications (Li *et al.*, 2008, 2013, 2016). In brief, the catheter of the probe was inserted into the descending aorta of the rat via the femoral artery for ABP. For EEG activity, two wire leads of HD-X11 transmitter were tunneled subcutaneously on to the surface of the parietal bone in the skull for animals at P7-30. For animals at P35-40, three EEG electrodes were screwed onto the skull and two EMG electrodes were sutured onto the dorsal neck muscles. All electrode leads were inserted into a sterilized six-prong plastic pedestal that was secured to the skull.

#### BrdU (Bromodeoxyuridine) injection

Newly proliferated OX neurons in the hypothalamus were marked using BrdU and identified by BrdU and OX-ir double staining. Rat pups were injected with BrdU (i.p. 50 mg kg^-1^ day^-1^ BrdU in saline; Roche Diagnostics, Indianapolis, IN, USA) for 5 consecutive days, i.e. from P5 to P9, P10 to 14, or P15 to P19, respectively (hereafter, the simplified P5-9, P10-14 and P15-19 injected rats terminology will be used). *Hcrt-SAP injection.* Elimination of excess OX neurons was achieved using a specific toxin for OX neurons, Hcrt-SAP (hypocretin-2-saporin, Advanced Targeting Systems, San Diego, CA, USA). The method of stereotaxic injection was similar to those described in our previous study and will be brief here (Nattie *et al.*, 2004; Li & Nattie, 2006). Rats were anesthetized with a ketamine (Putney, Inc., Portland, ME, USA) and xylazine (Lloyd Labs, Walnut, CA, USA) cocktail (90/15 mg kg^-1^, I.M.), and then fixed in a Kopf stereotaxic frame. Hcrt-SAP or its control IgG-SAP was unilateral injected (0.5 uL, 90 ng/ uL) into the hypothalamus with the coordinates at 1.4 mm lateral, 2.3 mm caudal from Bregma and 8.8 mm below the surface of the skull. All injections were made using a 1 uL Hamilton syringe, and each microinjection lasted at least 5 min and the needle remained in position for another 5 min before removal.

### Blood pressure and ventilatory measurements and data collection

The methods used to measure ABP, body temperature, and EEG were those in common use in our laboratory (Cummings *et al.*, 2011; Penatti *et al.*, 2011; Li *et al.*, 2013,2016). *For study in P15-P30 rats*, pups were placed in a water-jacketed glass chamber with body temperature held at 36 ± 0.5°C throughout the experiment by controlling the temperature of the water perfused through the chamber. The signals of ABP, core body temperature, EEG, and barometric pressure of HD-X11 telemetric probe were collected. *For study in P37-40 rats*, animals were placed in a traditional whole-body plethysmograph with food ad libidum as described previously (Li *et al.*, 2013*a*, 2016). Body temperature was measured before and after recording. Raw EEG and EMG outputs from the skull and neck skeletal muscle electrodes were filtered at 0.3–70 and 0.1–100 Hz, respectively, using a Grass Physiodata Amplifier System (NatusNeurology Inc., Grass Products, Middleton, WI, USA). All data were collected continuously throughout the experiment via a PhysioTel Connect device enabler system (DSI, St. Paul, MN, USA) using LabChart 8 software (ADInstruments, Colorado Springs, CO, USA). ABP signal was sampled at 1,000 Hz whereas breathing, EEG, and body temperature were sampled at 150 Hz. Heart rate (HR) was derived from pulse pressure of ABP.

### Histology

#### Tissue processing and harvesting

Rats were deeply anaesthetized with ketamine and xylazine and then transcardially perfused with saline followed by chilled 4% paraformaldehyde (PFA, 4% in 0.1 m phosphate buffer, pH 7.4). The brain was harvested and post-fixed overnight in 4% PFA at 4°C, then cryoprotected in 30% sucrose for 48 h. 40 *μ*m thick brain sections were used for immunohistochemical staining.

#### Immunohistochemical staining

The general methods used for all immunohistochemical staining procedures were similar to those described in our previous studies (Li *et al.*, 2016), and will be brief here. Free-floating sections were first incubated in a primary antibody at 4°C for 48 h followed by a secondary antibody at 4°C for 24 h or at room temperature (RT) for 2 h. Phosphate buffered saline (PBS) or 0.1% Triton X-100 PBS (PBST) were used for all washes between antibody incubations. Peroxidase and diaminobenzidine (DAB) with or without nickel were used for visualization.

*For OX-only staining*, brain sections were incubated in anti-orexin-A primary antibody (1:10,000 dilution; goat polyclonal, SC-8070, Santa Cruz, Dallas, TX, USA), followed by biotinylated horse anti-goat IgG secondary antibody (1:1,00 dilution; Vector laboratories, Burlingame, CA, USA). DAB with nickel was used for visualization (black neurons). *For BrdU and OX double staining*, a special treatment was used to enhance a better staining of BrdU. The brain sections were first denatured with 2N HCl for 45 minutes, followed by neutralization in 0.1M boric buffer for 30 minutes (min), and then washed three times with 0.1% Triton X-100 PBS (PBST). The sections were incubated in mouse anti-BrdU primary antibody (1:100 dilution; G3G4, Developmental Studies Hybridoma Bank, Iowa City, Iowa, USA) followed by a biotinylated horse anti-mouse IgG secondary antibody (1:1,000 dilution; Vector Laboratories, Burlingame, CA, USA). Peroxidase and diaminobenzidine with nickel were used to visualize the BrdU (black nuclei). Brain sections were then incubated in a goat polyclonal anti-OXA primary antibody (1:10,000 dilution; SC-8070, Santa Cruz, Dallas, TX, USA) followed by biotinylated horse anti-goat IgG secondary antibody (1:1,000 dilution; Vector Laboratories, Burlingame, CA, USA). Peroxidase and diaminobenzidine without nickel were used to visualize OX (brown neurons). Three types of neurons were identified for the quantification in this study (Figure 2B and C). BrdU/OX-ir neurons, positive for both BrdU and OX, were identified as brown neurons with black nuclei, which were likely postnatally proliferated OX neurons. OX-ir only neurons, positive for OX-ir but negative for BrdU, were identified as brown without black nuclei, which were generated prior to the postnatal period in this study. BrdU-ir only neurons, positive for BrdU but negative for OX-ir, identified as black nuclei only, which were likely postnatally generated non-OX neurons in the hypothalamus.

For cell counts, the hypothalamus was divided into three zones, the dorsomedial hypothalamus (DMH), the perifornical hypothalamus (PeF) and the lateral hypothalamus (LHA) in each hemisphere as described in the lab’s previous study (Li *et al.*, 2016), and all cell counts were completed using Neurolucida and Stereo Investigator (MFB Bioscience, Williston, VT, USA).

**Figure 1.**
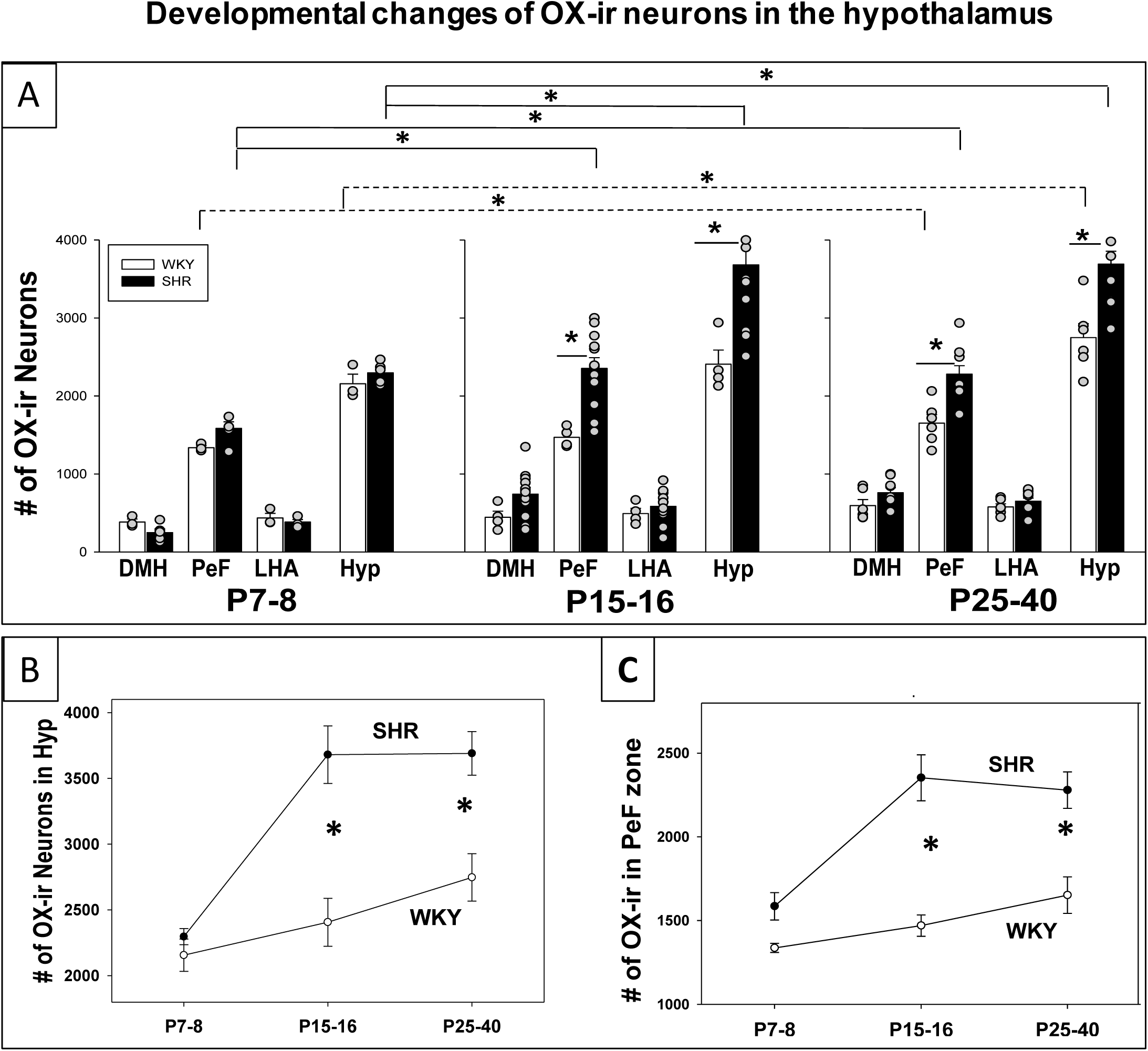
Postnatal developmental changes of OX neurons in the hypothalamus in SHRs and WKY rats. Total number of OX neurons in the hypothalamus (Hyp) and three hypothalamic zones (DMH, LHA, PeF) at three developmental ages, P7-8, P15-16, and P25-40 in SHRs and WKY rats are shown in A. *Indicates significance, Two-way ANOVA with Holm-Sidak or Student-Newman-Keuls *post hoc* tests. B-C show the comparison of OX neurons in the hypothalamus and PeF zone between SHRs and WKY rats at three developmental ages. *Indicates significance between SHRs and WKY rats. Abbreviations: DMH, dorsomedial hypothalamus; PeF, perifornical zone; LHA, lateral hypothalamic area. Data are shown as mean ± SEM.

**Figure 2.**
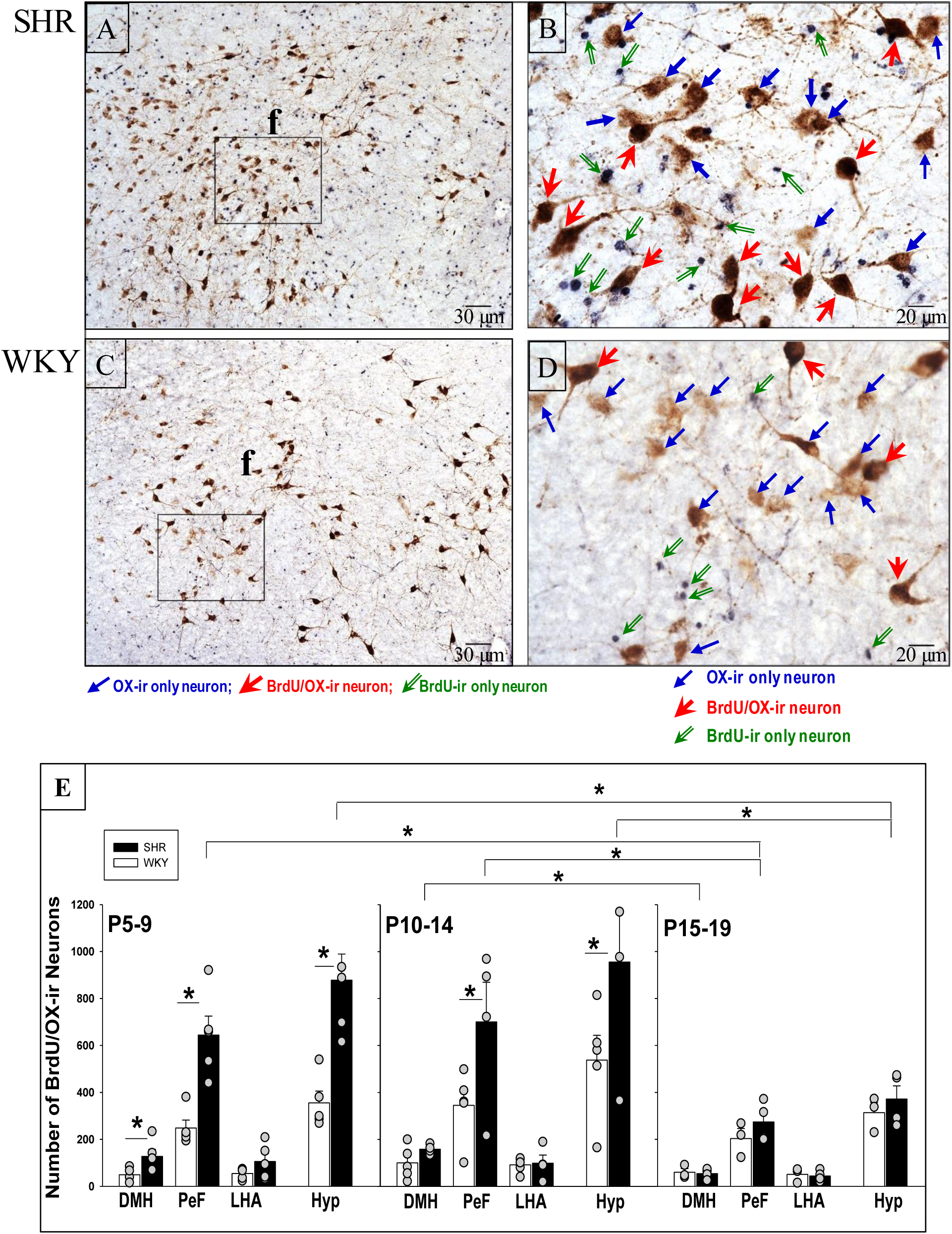
Postnatal OX neurogenesis in SHRs and WKY rats at three developmental periods. Representative images show OX-ir and BrdU/OX-ir neurons within comparable hemispheres of the hypothalamus in SHRs and WKY rats from P5-9 group (A-D), and expanded view of OX-ir neurons (blue arrows; brown neurons), BrdU/OX-ir neurons (red arrows; brown neurons with black nuclei), and BrdU-ir neurons (green arrows; black nuclei), (B, D). E shows the total number of BrdU/OX-ir neurons in the Hyp and the distribution within the three zones in SHRs and WKY rats in each BrdU-injected age group. *Values are significantly different based on Bonferroni *post hoc* test. Abbreviations: DMH, dorsomedial hypothalamus; PeF, perifornical zone; LHA, lateral hypothalamic area; BrdU, bromodeoxyuridine; OX, orexin-A; ir, immunoreactive. Data are shown as mean ± SEM.

### Experimental designs and data analysis

#### Experiment 1: *Postnatal developmental changes in number of orexin neurons in SHRs*

To determine at which developmental age the number of OX-ir neurons in SHRs diverges from normotensive WKY rats. SHR and WKY rat pups were divided into three age groups, P7-8 (SHR n=5; WKY n=4), P15-16 (SHR n=12; WKY n=4), and P25-40 (SHR n=10; WKY n=8). Brains were harvested for quantification of OX-ir neuron at three postnatal developmental ages.

#### Data analysis

The total number of OX-ir neurons were quantified and compared between SHRs and WKY rats at three ages (P7-8, P15-17, and P25-40) using a two-way ANOVA with age and strain (SHR and WKY rats) as the factors. A Holm-Sidak or Student-Newman-Keuls *post hoc* test was applied when appropriate. All values are reported as mean ± SEM.

### Experiment 2: *Postnatal neurogenesis of orexin neurons*

To determine whether SHR pups have more postnatal proliferation of orexin neurons than age matched normotensive WKY pups during development. The BrdU injected rats were divided into three age groups P5-9 (SHR n=5; WKY n=3), P10-14 (SHR n=3; WKY rats n=4), and P15-19 (SHR n=4; WKY rats n=3) with similar numbers of males and females in each group. The brains were harvested 1 day after the 5^th^ injection of BrdU except for P5-9 injected rats, which were harvested 3-5 days after the conclusion of BrdU injections.

#### Data analysis

The total number of OX neurons that are positive for BrdU (BrdU/OX-ir) in the hypothalamus were quantified and compared between SHRs and WKY rats at three age groups using a two-way ANOVA with age and strain (SHR and WKY rats) as the factors. Additionally, the differences in the number of BrdU/OX-ir in three hypothalamic zones, DMH, PeF and LHA, were compared between SHRs and WKY rats at P5-9, P10-14 and P15-19 using a two-way ANOVA with age and strain (SHR and WKY) as the factors. A *post hoc* Holm-Sidak multiple comparison was applied when appropriate. All values are reported as mean ± SEM.

### Experiment 3: *Chronological relationship between development of excess OX neurons and higher ABP in SHRs*

To determine at which developmental age SHRs begin to have a significantly higher ABP than age matched normotensive WKY controls and its chronological relationship with the postnatal changes in the OX system. Rat pups at four ages were used to measure the developmental change of ABP, P15 (SHR n=5; WKY n=4), P20 (SHR n=9; WKY n=6), P25 (SHR n=8; WKY n=8), and P30 (SHR n=11; WKY n=8) (table 1). One day after implantation of BP telemetry and EEG, rat pups were acclimatized in a water-jacketed chamber for at least 1 hour and body temperature was maintained at 36 ± 0.5°C. ABP, EEG, and body temperature in wakefulness and sleep were collected continuously for 1.5-3 hours in room air conditions.

**Table 1.** Body weight at each age in experimental 3.

|  | **P15** | **P20** | **P25** | **P30** |
| --- | --- | --- | --- | --- |
| **SHR** | 22.91±1.46 g | 34.72±8.11 g | 41.76±8.87 g | 59.34±11.22 g |
| **WKY** | 32.34±3.61 g | 33.37±3.63 g | 40.20±7.87 g | 56.20±9.81 g |

Body weight at P15, P20, P25 and P30 of SHR and WKY rat pups used in Experiment 3.
Both SHR and WKY rat pups had age-dependent increase in body weight ( $P \leq 0.001$ ).

#### Data Analysis

The mean ABP of SHR and WKY rat pups in wakefulness and NREM sleep at P15, P20, P25, and P30 were analyzed and compared using a two-way ANOVA with age and strain (SHR and WKY rats) as the factors. A Holm-Sidak or Student-Newman-Keuls *post hoc* test was applied when appropriate. All values are reported as mean ± SEM. Chronological relationship between development of excess OX neurons and higher ABP in SHRs and WKY rats is shown in Figure 3B.

**Figure 3.**
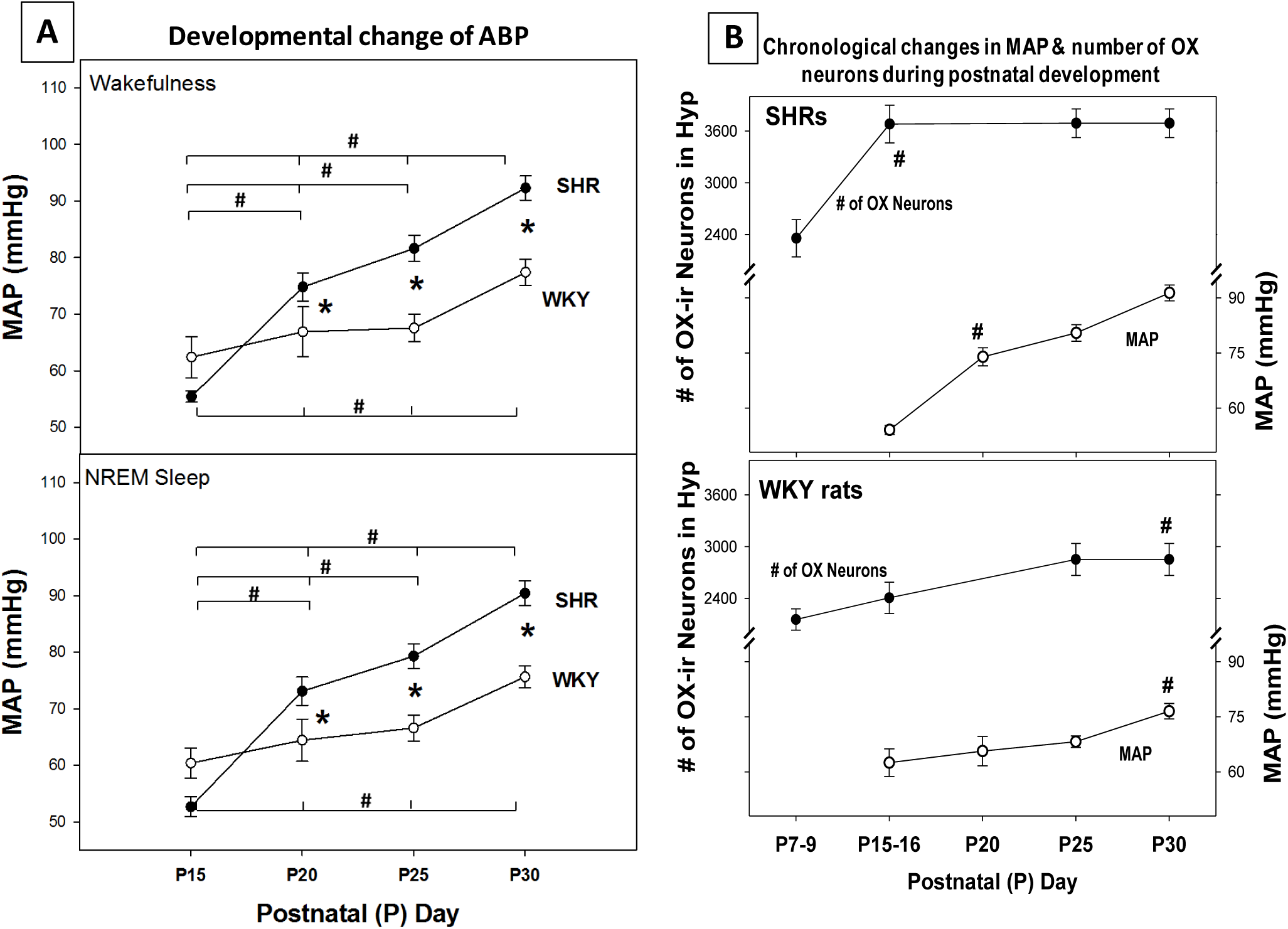
Postnatal changes of mean arterial blood pressure (MAP) in SHRs and WKY rats. Panel A shows MAP at P15, P20, P25 and P30 in SHR and normotensive WKY rats in wakefulness and sleep. Chronological relationship between MAP (right y-axis) and number of OX neurons (left y-axis) in SHR and WKY rats during postnatal development is shown in B. *p<0.05, values are significantly different between SHR and WKY rat pups; #p<0.05, values are significantly different with age in SHR or WKY rat. Two-way ANOVA with Holm-Sidak *post-hoc* test. Data are shown as mean ± SEM.

### Experiment 4: *The effects of eliminating excess orexin neurons on ABP and ventilatory chemoreflex*

To determine whether eliminating excess orexin neurons via Hcrt-SAP can prevent SHRs from developing a higher ABP and exaggerated hypercapnic response two groups of young SHRs (P30-40) were used. Ages of both groups at the day of injection and at the day of physiological experiments are included in Table 2. SHRs were randomly assigned into two groups to receive either Hcrt-SAP or IgG-SAP injection between P25-28 (n=6, WT 90.4g; n=5, WT 82.2g, respectively). Twelve days post-injection at P37-40, SHRs were allowed to acclimatize for 1-2 hours in an experimental whole-body plethysmography. ABP and ventilatory data in wakefulness and sleep were collected for 1-2 hours during the light period in room air, 1-2 hours during the dark period in room air, and 1-2 hours during the dark period in a 5% CO_2_ mixed gas (5% CO_2_ 21% O_2_ balanced with nitrogen). At conclusion of the recording, the brain was harvested for OX-ir staining and quantification.

**Table 2.** Age and body weight of SHRs used in experiment 4.

| **Treatment group** | **Number of animals** | **Age at injection** | **BW at injection (g)** | **Age at physiology experiment** | **BW at physiology experiment (g)** |
| --- | --- | --- | --- | --- | --- |
| Hcrt-SAP | 6 | 26.2 ± 0.5 | 56.1 ± 2.6 | 38.2 ± 0.5 | 90.4 ± 4.4 |
| IgG-SAP | 5 | 27.2 ± 0.6 | 52.7 ± 3.6 | 39.2 ± 0.6 | 82.2 ± 5.8 |

There was no statistically significant differences in body weight or age at the day of injection or the day of physiological measurement between Hcrt-SAP and IgG-SAP groups. Data are shown as mean ± SEM. Abbreviations: BW, body weight.

#### Data Analysis

The number of OX neurons, mean ABP and ventilation were compared between Hcrt-SAP and IgG-SAP treated SHRs 12 days post injection. Two levels of analysis were utilized to evaluate the extent of elimination of orexin neuron induced by Hcrt-SAP. First, to determine the difference in the total number of OX-ir neurons between injected hemisphere and non-injected hemisphere in the hypothalamus and the three individual zones in same SHRs treated with either IgG-SAP or Hcrt-SAP (Fig 4-C) a One-Way ANOVA was used to compare the difference between two hemispheres. Second, to determine the total loss of OX neurons induced by Hcrt-SAP the difference in the percentage loss of OX neurons in the injected hemisphere between Hcrt-SAP and IgG-SAP treated rats a two-way ANOVA was used with treatments and regions as factors (Fig 4-B). Holm-Sidak *post hoc* for multiple comparisons was used when appropriate. For the physiological effects, mean ABP and ventilation were compared between Hcrt-SAP and IgG-SAP treated animals within the respected measuring light period using a two-way ANOVA analysis with treatment and vigilance state as factors (Fig 5). Holm-Sidak *post hoc* for multiple comparisons when appropriate All values are reported as mean ± SEM.

**Figure 4.**
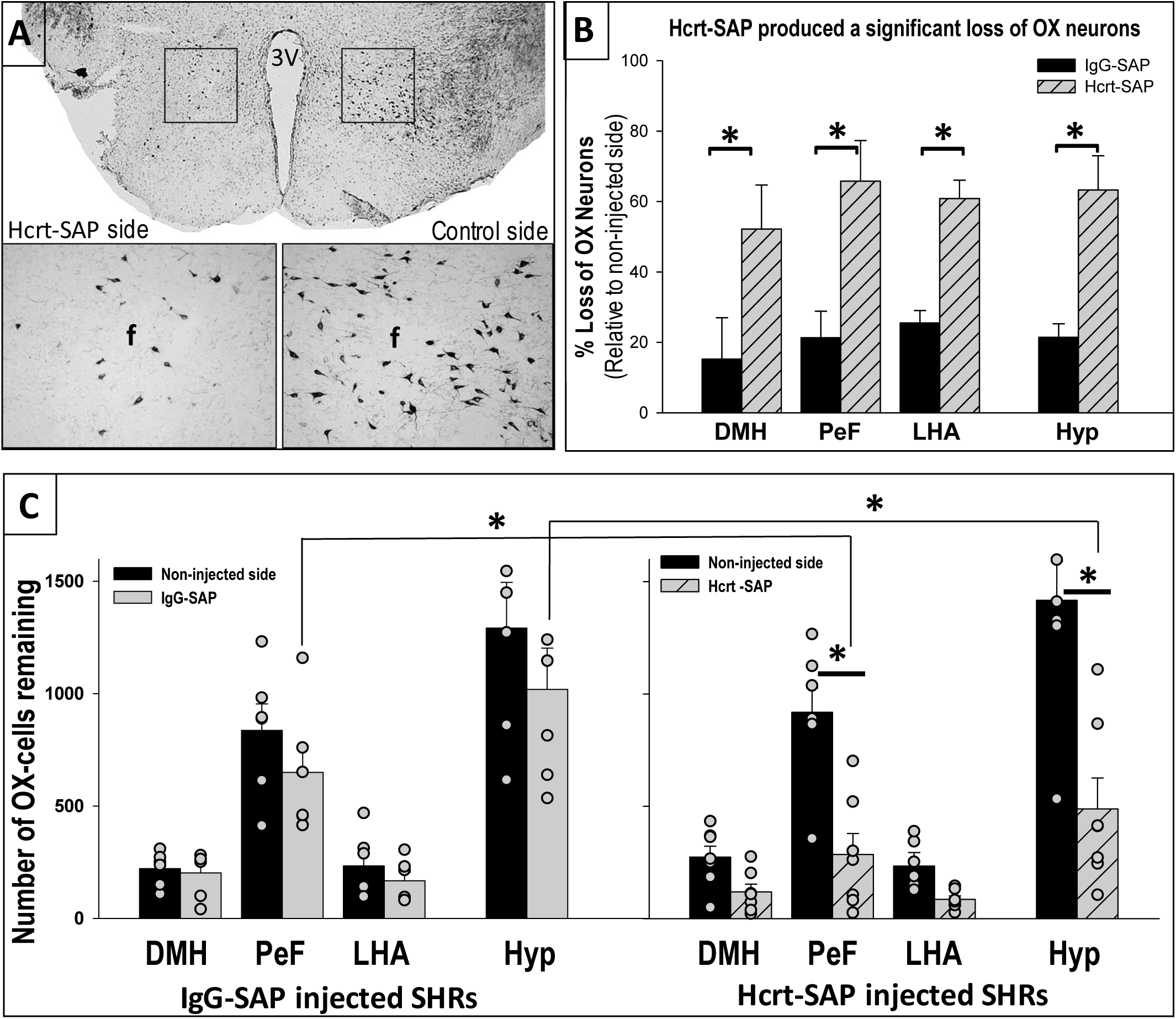
Verification of Hcrt-SAP effect on eliminating excess OX neurons in SHRs. Representative images (A) top panel show a full view of the hypothalamus (10X) of Hcrt-SAP injected (left) and non-injected (right) hemisphere, and lower panels show higher magnification (20X) photomicrographs of the areas encompassed by the squares of top panel, lesion (left) vs control (right) sides. 4B shows a comparison of percentage loss of OX neurons in the injected hemisphere relative to the non-injected hemisphere between Hcrt-SAP and IgG-SAP injected SHRs. 4C shows comparisons of total number of OX neurons between non-injected (black bar) and injected (grey bar) hemisphere in Hcrt-SAP and IgG-SAP injected SHRs, and in the injected hemisphere between Hcrt-SAP (hatched bars) and IgG-SAP injected SHRs. *p<0.05, Two-way ANOVA with Holm-Sidak *post-hoc* test. Abbreviations: hcrt-SAP, hypocretin-2-saporin; IgG-SAP, IgG-saporin; OX, orexin-A; f, fornix; DMH, dorsomedial hypothalamus; PeF, perifornical zone; LHA, lateral hypothalamic area; Hyp, total hypothalamus; SHR, spontaneously hypertensive rats. Data are shown as mean ± SEM.

**Figure 5.**
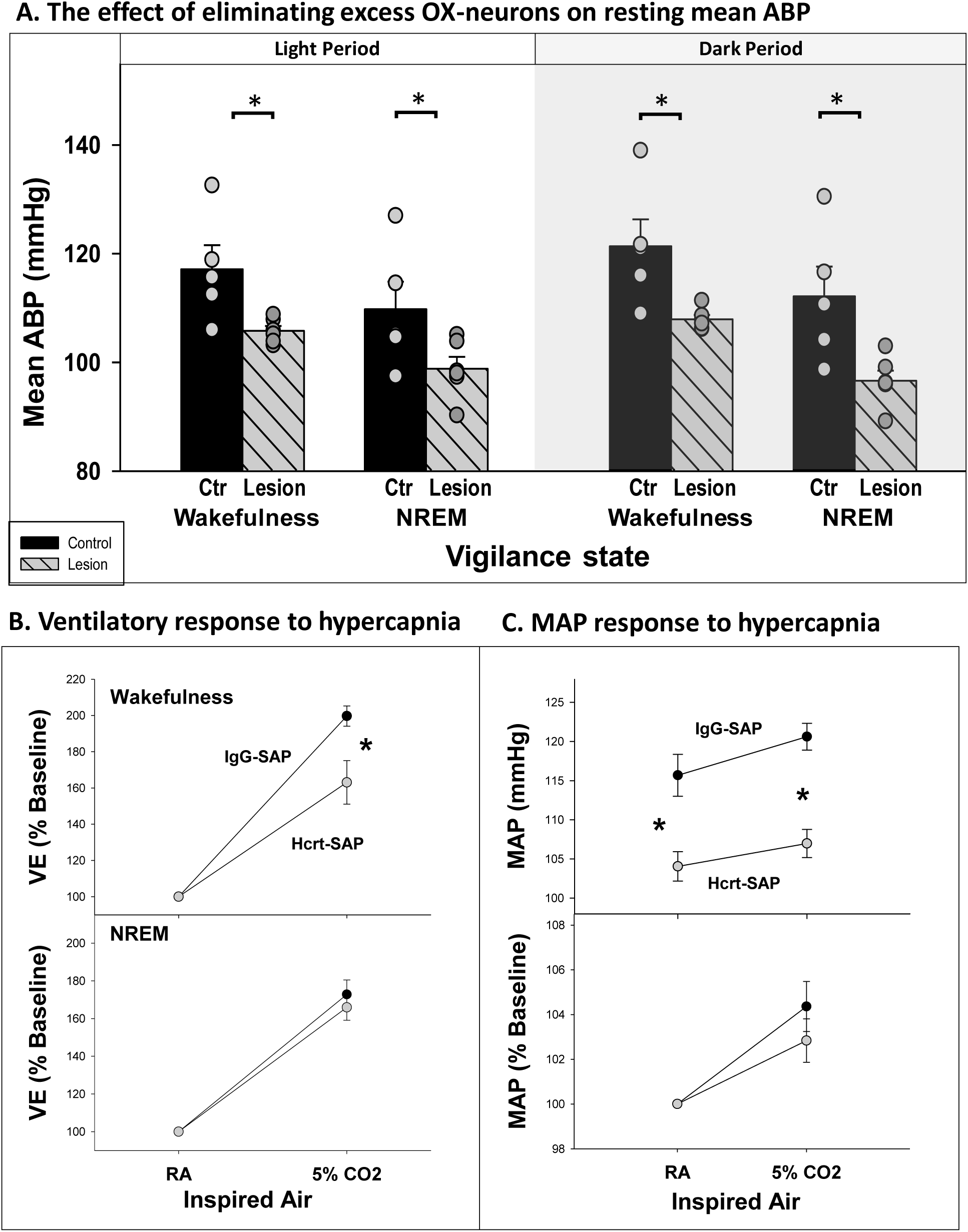
The effects of elimination of excess OX neurons on ABP and CO_2_ chemoreflex in SHRs at a developmental period. A comparison of the effects on resting mean ABP between Hcrt-SAP and IgG-SAP injected SHRs in quiet wakefulness and NREM sleep during the light and dark cycle is shown (A). Hcrt-SAP injected SHRs had a significantly lower ABP than IgG-SAP injected SHRs during the dark cycle in both NREM sleep and quiet wakefulness. Ventilatory response to hypercapnia (B) was significantly lower in Hcrt-SAP injected SHRs in wakefulness. ABP response to hypercapnia was not different between Hcrt-SAP and IgG-SAP injected SHRs despite Hcrt-SAP injected SHRs have significantly lower resting ABP (C). *p<0.05, Two-Way ANOVA with Holm-Sidak *post-hoc* test. Data are shown as mean ± SEM. Abbreviations: NREM, non-rapid eye movement sleep; Ctr, controls; Hcrt-SAP, hypocretin-2-saporin; IgG-SAP, IgG-saporin; VE, ventilation; MAP, mean arterial blood pressure; RA, room air.

## Results

### Experiment 1: *Postnatal developmental changes in number of orexin neurons in SHRs*

The number of OX-ir neurons in the three hypothalamic zone (DMH, PeF, and LHA) and the hypothalamus (Hyp; three zones combined) at three developmental ages (P7-8, P15-16 and P25-40) are summarized and shown in Figure 1. Both SHRs and WKY rats experienced developmental increases in the number of OX-ir neurons in the hypothalamus, however SHRs started to have significantly more OX neurons from P15 onward (Figure 1).

In SHR pups, there was a significant postnatal surge in the total number of OX-ir neurons in the Hyp during postnatal development. SHR pups at ages P15-16 (3680 ± 219) and P25-40 (3690 ± 166) had significantly more OX-ir neurons in Hyp than the pups at P7-8 (2359±216) (P<0.001; Two-way ANOVA with Holm-Sidak analysis; Figure 1). The increase in total number of OX neurons in the Hyp was primarily contributed by the PeF zone, where significantly more OX-ir neurons were found in pups at P15-16 and P25-40 than at P7-8 (P<0.001; Two-way ANOVA with Holm-Sidak analysis; Figure 1 B-C). Additionally, a similar difference, but on a much smaller scale, was also found in both the DMH and LHA zones where SHR pups at P15-16 (P<0.001 & P=0.017, respectively) and P25-40 (P<0.001 & P=0.02, respectively) had more OX-ir neurons than the pups at P7-8 (Two-way ANOVA with Holm-Sidak analysis; Figure 1 A).

In WKY rats, there was a noticeable small increase in total number of OX-ir neurons in the hypothalamus with age, however the change was not statistically significant among the three age groups (P7-8, P15-16 and P25-40) (P>0.05; Two-way ANOVA with Holm-Sidak analysis, Figure 1). Further analysis showed that there was a small but significant increase in the number of OX-ir neurons in the PeF zone with age, where WKY rat pups at ages P25-40 (1687 ± 102) had significantly more OX-ir neurons than the pups at P7-8 (1337±27) and P15-16 (1470 ± 64) (P=0.005 & P<0.001, respectively, Figure 1C). There was no significant difference in the number of OX-ir neurons in the LHA and DMH zones at any age in WKY rat pups (P>0.05; Two-way ANOVA with Holm-Sidak analysis; Figure 1 A-B).

Between SHR and WKY pups, there was an age-related difference in the number of OX-ir neurons in the hypothalamus, particularly in the PeF zone, during postnatal development between P15-P40 (P<0.001; Two-way ANOVA with Holm-Sidak analysis; Figure 1 A-C). At P7-8, there was no statistical difference in the number of OX-ir neurons in the Hyp (2359 ± 216 vs 2156 ± 123, respectively) or any of the three hypothalamic zones (DMH, PeF, and LHA) between SHR and WKY rat pups (P>0.05; Two-way ANOVA with Holm-Sidak analysis; Figure 1 A-C). At P15-16, a significant difference in total number of OX-ir neurons in the Hyp between SHRs (3680 ± 219) and WKY rats (2407 ± 182) emerged, and the difference was primarily contributed by the PeF zone (2423 ± 144 vs 1470 ± 64, respectively), (P<0.001; Two-way ANOVA with Holm-Sidak analysis, Figure 1). Additionally, there was a small but significant difference in the number of OX-ir neurons in the DMH zone between SHR (742±85) and WKY (445±77) rat pups at P15-16 (P=0.023; Two-way ANOVA with Holm-Sidak analysis; Figure 1 A). Such differences between SHR and WKY rat pups persisted into ages P25-40 in the Hyp (3690 ± 166 vs 2852 ± 185, respectively) and in the PeF zone (2242 ± 96 vs. 1687 ± 102, respectively), (P<0.001; Two-way ANOVA with Holm-Sidak analysis; Figure 1 A-B). There were no age-related significant differences in the number of OX-ir neurons in the LHA zone between SHR and WKY rat pups during the postnatal developmental periods studied (P>0.05; Two-way ANOVA with Holm-Sidak analysis; Figure 1 A).

### Experiment 2: *Postnatal neurogenesis of orexin neurons*

The representative images of newly proliferated OX-ir neurons marked by BrdU and the number of BrdU/OX-ir neurons in the three hypothalamic zones (DMH, PeF, and LHA) and the hypothalamus (Hyp; three zones combined) at the three developmental ages studied (P5-9, P10-14 and P15-19) in SHR and WKY pups are shown in Figure 2.

In SHRs, there were age-related differences in the number of newly proliferated BrdU/OX-ir neurons in the hypothalamus, P5-9 (878 ±112) and P10-14 (956 ± 208) BrdU injected pups had significantly more BrdU/OX-ir neurons in the hypothalamus than P15-19 injected pups (372 ± 56) (P=0.004 and 0.002, respectively; Two-way ANOVA (age and region) with Holm-Sidak analysis; Figure 2 E). The majority of BrdU/OX-ir neurons were found in the PeF zone, where significantly more BrdU/OX-ir neurons were observed in P5-9 (644 ±181) and P10-14 (701±169) injected pups than in P15-19 injected pups (274 ± 42), (P=0.002 & 0.007, respectively; Two-way ANOVA (age and region) with Holm-Sidak analysis; Figure 2 E). It is also notable that there was a small but significant difference in the number of BrdU/OX-ir neurons in the DMH zone between P10-14 (159±10) and P15-19 (54±11) injected SHR pups (P=0.005; Two-way ANOVA with Holm-Sidak analysis; Figure 2 E).

In WKY rats, P10-14 BrdU injected pups appeared to have more BrdU/OX-ir neurons in the PeF zone and Hyp (345±66 & 538±106, respectively) than P5-9 (248±50 & 355±50, respectively) and P15-19 (203±42 & 314±43, respectively) injected pups, but differences were not statistically significant (P>0.05; Two-way ANOVA with Holm-Sidak analysis; Figure 2 E).

When comparing the difference between SHR and WKY rat pups, there were age-specific differences in the number of newly proliferated BrdU/OX-ir neurons in the hypothalamus (P≤0.01; Two-way ANOVA with Holm-Sidak analysis, age and strain as factors; Figure 2 E). P5-9 (878±112) and P10-14 (956±208) SHR pups had significantly more BrdU/OX-ir neurons in the hypothalamus than that of age matched WKY rat pups (355±50 & 538±106, respectively), (P=0.02 & P=0.015, respectively; Two-way ANOVA with Holm-Sidak analysis; Figure 2 E). The primary difference in the number of newly proliferated orexin neurons was in the hypothalamic PeF zone, where P5-9 (644±181) and P10-14 (701±169) injected SHR pups had significantly more BrdU/OX-ir neurons than that of age-matched WKY rat pups (248±50 & 345±66, respectively), (P=0.002 & 0.007, respectively; Two-way ANOVA with Holm-Sidak analysis; Figure 2 E). There was also a small but significant difference in the number of BrdU/OX-ir neurons in the DMH zone between P5-9 injected SHR (127±30) and WKY rat (49±12) pups (P=0.017; Two-way ANOVA with Holm-Sidak analysis). There was no difference in the number of BrdU/OX-ir neurons in the Hyp or any of three hypothalamic zones between P15-19 BrdU injected SHR and WKY rat pups (P>0.05; Two-way ANOVA with Holm-Sidak analysis). No significant difference was observed in the LHA zone between SHR and WKY rat pups at any of the three ages studied (P>0.05; Two-way ANOVA with Holm-Sidak analysis; Figure 2 E).

It is worth noting that all brains from P5-9 BrdU injected SHR and WKY rats were harvested between P12-14, in which BrdU-ir marked neurons were generated between P5-9 while co-stained OXA marked OX neurons that were between P5-P14 old. Nevertheless, even when combining BrdU/OX-ir from both P5-9 and P10-14 injected rats there is clearly more newly proliferated OX (BrdU/OX-ir) neurons in the hypothalamus in P5-9 and P10-14 BrdU injected SHRs than age-matched WKY rats. BrdU is commonly used to identify newly proliferating cells in the brain (Miller & Nowakowski, 1988; Cameron & Mckay, 2001; Menyhárt *et al.*, 2016; Chang *et al.*, 2008, 2012, 2013), and multiple 50 mg kg^−1^ injections (i.p.) of BrdU can specifically and sufficiently label newly generated neurons (Wojtowicz & Kee, 2006) (Cameron & Mckay, 2001)(Miller & Nowakowski, 1988; Takahashi *et al.*, 1992).

### Experiment 3: *Chronological relationship between development of excess OX neurons and higher ABP in SHRs*

Developmental changes of ABP were accessed in SHRs and their normotensive controls at P15, P20, P25, and P30 to determine its chronological relationship with excess OX neurons during postnatal development (*Figure 3*). The general conditions are similar between age matched SHRs and their background normotensive control WKY pups. There were age-related increases in body weight in both SHRs and WKY rats, however there were no significant differences in body weight between the two strains at any age (P>0.05; Table 1). Mean ABP of SHR and WKY rat pups in wakefulness and sleep at P15, P20, P25 and P30 are shown in Figure 3.

Both SHRs and WKY rats experienced postnatal developmental increases in ABP (P<0.001; Two-way ANOVA, age and strain as factors; Figure 3). In SHR pups, an age-dependent increase in ABP was observed in both wakefulness and sleep among the age groups, P15 (55.4 ± 1.0 and 52.7 ± 1.7 mmHg, respectively), P20 (74.8 ± 2.5 and 73.1 ± 2.5 mmHg, respectively), P25 (81.6 ± 2.3 and 79.3 ± 2.2 mmHg, respectively), and P30 (92.3 ± 2.2 and 90.4 ± 2 mmHg, respectively), (P<0.001; Two-way ANOVA with Holm-Sidak analysis, Figure 3). In normotensive WKY rat pups, mean ABP also rose with age, and there were significant differences between age P30 and P15, P20, and P25 (P≤0.011), however the change was much smaller compared to age matched SHRs (Figure 3)(Two-way ANOVA with Holm-Sidak analysis).

Between SHR and WKY rat pups, there was an age-dependent difference in mean ABP during the developmental period studied, and there was a statistically significant interaction between age and strain (SHR vs WKY) in wakefulness (P=0.003) and sleep (P=0.002), (Two-way ANOVA with Holm-Sidak analysis, age and strain as factors). At P15, there was no significant difference in resting mean ABP in wakefulness and NREM sleep between SHRs (55.4 ± 1.0 & 52.7 ± 1.7 mmHg, respectively) and WKY rats (62.4 ± 3.6 & 61.6 ± 3.9 mmHg, respectively), (P≥0.05; Two-way ANOVA with Holm-Sidak analysis). At P20, a small but significant difference in mean ABP between SHR and WKY rat pups emerged, and mean ABP in wakefulness and sleep, respectively, was 74.8 ± 2.5 and 73.1 ± 2.5 mmHg in SHR *vs* 66.9 ± 4.4 and 62.4 ± 3.7 mmHg in WKY rats (P=0.042 and 0.017, respectively; Two-way ANOVA with Holm-Sidak analysis). The difference in mean ABP between SHR and WKY rat pups magnified with age, and mean ABP in wakefulness in SHR and WKY rat pups was 81.6 ± 2.3 vs 67.5 ± 2.4 mmHg, respectively, at P25, and 92.3 ± 2.2 vs 77.4 ± 2.3mmHg, respectively, at P30 (Figure 3; P<0.001; Two-way ANOVA with Holm-Sidak analysis). A similar age-dependent difference in mean ABP in NREM sleep between SHR and WKY rat pups was also observed (Figure 3A).

The chronological relationship between surges of OX neurons and ABP in SHRs *vs* age-matched WKY are shown in Figure 3-B. In SHRs, the surge in number of OX neurons emerged at P15, about 5 days prior to the aberrant rise in ABP, which suggests that increased OX activity may be necessary for developing a higher blood pressure in SHRs during postnatal development.

### Experiment 4: *The effects of eliminating excess orexin neurons on ABP and ventilatory chemoreflex*

To determine whether excess OX activity is necessary in developing hypertension during development we evaluated the effects of eliminating some OX-neurons via Hcrt-SAP on ABP and ventilatory hypercapnic chemoreflex in SHRs during a developmental period (Figures 4-5).

Ages and body weights at time of injection and at the time of the physiology experiment are recorded in Table 2. There were no significant differences in the age of injection between Hcrt-SAP and IgG-SAP injected SHRs (26.2 ± 0.5 vs 27.2 ± 0.6, respectively), body weight at injection (56.1 ± 2.6 vs 52.7 ± 3.6, respectively), age at time of physiology experiment (38.2 ± 0.5 vs 39.2 ± 0.6, respectively), or body weight at time of physiology experiment (90.4 ± 4.4 vs 82.2 ± 5.8, respectively). Physiology experiments were completed exactly 12-days post-lesion for each individual animal with injection ages ranging from P25-P28 and experimental ages ranging from P37-P40.

#### Efficacy of eliminating excess OX neurons

The representative images of Hcrt-SAP injected and non-injected hemispheres in a SHR are shown in Figure 4-A, and the quantified effect of Hcrt-SAP on number of OX neurons are shown in Figure 4B-C.

To determine the number of remaining OX neurons in the hypothalamus, the number of OX neurons were compared between the injected and non-injected hemisphere in the same SHR. In IgG-SAP injected control SHRs, there was a very small decrease in number of OX neurons in the injected hemisphere relative to the non-injected hemisphere, however the loss was not statistically significant (P>0.05, two-way ANOVA) and was likely due to mechanical injury during the injection procedure (Figure 4C left panel). While in Hcrt-SAP injected SHRs, there was a significant decrease in total number of OX neurons resulting from the Hcrt-SAP lesion in injected hemisphere vs non-injected hemisphere in Hyp (∼488 *vs* 1417, respectively) and PeF zone (285 *vs* 918, respectively) (P≤0.001, two-way ANOVA with treatment and region as factors; Figure 4C, right panel).

When comparing the cell loss in the injected hemisphere *between Hcrt-SAP and IgG-SAP injected SHRs* the percentage loss of OX neurons was calculated by dividing the total number of remaining OX-ir neurons in the injected hemisphere over the total number of OX-ir neurons in the non-injected hemisphere. In the injected hemisphere, Hcrt-SAP injected SHRs lost significantly more OX neurons than IgG-SAP injected control SHRs in the hypothalamus (63.3±10 % vs 21.4 ±4%, respectively, P=0.002), and three hypothalamic zones, DMH, PeF, and LHA (52.2% vs 15.2%; 65.8% vs 21.3%; and 60.8% vs 25.5%, respectively, P≤0.05, Two-way ANOVA with Holm-Sidak multiple comparisons, *Figure 4B).* Additionally, we estimated the total loss of OX neurons in the whole brain based on the assumption that both hemispheres have an equal number of OX neurons if there was no injection. Thus, the total percentage loss of OX neurons in the whole brain would be equal to ½ of the percent loss of OX in the injected hemisphere mentioned above. In Hcrt-SAP injected SHRs, the estimated total percent loss of OX neurons in the brain was 31.6%, which is half of percentage loss of injected hemisphere (63%), verses 10.7% in IgG-SAP control SHRs (P=0.002, two-way ANOVA with *Holm-Sidak multiple* comparisons). This number is relevant, as previous studies have shown that SHRs have ∼ 30% more OX neurons than age-matched normotensive controls.

#### The effects of eliminating excess OX neurons on resting ABP

The effects of elimination of excess OX neurons on resting ABP and both ventilatory and ABP response to hypercapnic challenge are shown in Figure 5. During the dark period in room air condition, Hcrt-SAP lesioned SHRs had a significantly lowered mean ABP compared to IgG-SAP control SHRs in NREM sleep (97±2 vs 112±6, respectively; P=0.006) and QW (108 ± 1 vs 122 ± 5, respectively; p=0.015)(Two-way ANOVA with Holm-Sidak multiple comparisons; Figure 5A). Similar results were found in the light period where Hcrt-SAP lesioned SHRs also had a lower mean ABP than IgG-SAP control SHRs in NREM sleep (98.9 ± 2.2 vs 109.8 ± 5.1, respectively; P=0.031) and QW (105.8 ± 0.9 vs 117.1 ± 4.4, respectively; P=0.026), (Two-way ANOWA with Holm-Sidak multiple comparisons; Figure 5A).

#### The effects of eliminating excess OX neurons on ABP and ventilatory responses to hypercapnia

The ABP response to hypercapnia (5%CO_2_) describes the ABP in the first 5 minutes after CO_2_ reached 4%. Mean ABP was significantly lower in Hcrt-SAP lesioned SHRs than IgG-SAP control SHRs in QW (107.9 ± 1.9 vs 120.6 ± 1.7, respectively; P<0.01, mixed-effects analysis with Sidak’s multiple comparisons) in both resting (room-air) and hypercapnic (5%CO_2_) conditions (Figure 5C). However, there was no significant difference in the change of ABP in response to CO_2_ between lesioned and control SHRs (Figure 5C). In terms of the ventilatory response to hypercapnia, Hcrt-SAP lesioned SHRs had a significantly lower ventilatory response to hypercapnia (5% CO_2_) than IgG-injected control SHRs in QW (61.6 ± 11.5 vs 89.5 ± 4.5, respectively, P<0.05 mixed-effects analysis with Sidak’s multiple comparisons, Figure 5), but not in NREM sleep (50.6 ± 6.0 vs 59.4 ± 6.2, respectively). The decreased CO_2_ chemoreflex was primarily contributed by a change of tidal volume (data not shown).

## Discussion

An overactive orexin system has been recently linked to neurogenic hypertension in adult SHRs (Lee *et al.*, 2013; Li *et al.*, 2013, 2016), however, the existence of a chronological relationship between the increased orexin activity and ABP and exaggerated OX neurogenesis during postnatal development was unknown until now. Here, we demonstrated that 1) postnatal orexin activity, marked by the number of OX-peptide positive neurons, surged to an abnormally high level in hypertensive rats compared to age-matched normotensive WKY controls by the age of P15, which was about 5 days prior to a measurable divergence of mean ABP between SHRs and WKY rats at ∼P20; 2) exaggerated postnatal orexin neurogenesis was the primary contributor to the excess OX neurons in the hypothalamus during postnatal development; and 3) eliminating excess OX neurons in the hypothalamus can prevent the development of a higher ABP and exaggerated hypercapnic chemoreflex in SHRs during a postnatal period (P25-40). The fact that a surge in the number of OX neurons occurs before an aberrant increase in mean ABP and that elimination of excess OX neurons can prevent developing a higher ABP during a postnatal developmental period suggests that the orexin system may play an important role in the development of neurogenic hypertension. We further suggest that an overactive orexin system may be necessary in developing neurogenic hypertension in SHRs, and modulation of such an overactive system may be beneficial in treating hypertension.

### Postnatal development of excess OX neurons via exaggerated OX neurogenesis in SHRs

Development and maturation of the orexin system occurs during late embryotic and early postnatal periods in normal rodents (Van den Pol *et al.*, 2001; Steininger *et al.*, 2004; Amiot *et al.*, 2005; Sawai *et al.*, 2010; Stoyanova *et al.*, 2010; Iwasa *et al.*, 2015). Early *in situ* hybridization studies showed that *prepro-orexin* mRNA expression is first seen between E12-18 (Steininger *et al.*, 2004; Amiot *et al.*, 2005) and the levels of expression progressively increase until about P30, at which age it remains stable into adulthood (Yamamoto *et al.*, 2000; Iwasa *et al.*, 2015). The number of orexin neurons, marked by the expression of either *prepro-orexin* mRNA or orexin-A and/or –B peptide, progressively increases in normal animals during postnatal development (Steininger *et al.*, 2004; Sawai *et al.*, 2010; Ogawa *et al.*, 2017). In adult hypertensive rodents, it has been previously reported that spontaneously hypertensive rats and mice have about 30 and 20% more OX-ir neurons, respectively, than their age-matched normotensive controls (Clifford *et al.*, 2015; Jackson *et al.*, 2016; Li *et al.*, 2016). We further reported that such increase in orexin activity can be observed in a pre-hypertension period at P30-58 in young SHRs (Li *et al.*, 2016). However, it was not clear whether SHRs were born with excess OX neurons or if the excess population developed postnatally.

In this study, we found that SHRs and normotensive WKY rats had similar numbers of OX neurons at P7-8, however by P15, SHRs started to have significantly more OX neurons than age-matched normotensive controls (Figure 1). These data suggest that SHR pups experienced an exaggerated surge in the total number of OX neurons postnatally between P7 and P16, which is primarily contributed by increased OX neurogenesis. Compared to age-matched normotensive WKY pups, SHR pups had significantly more newly proliferated OX neurons, which were positive for both BrdU and OXA (BrdU/OX-ir), in the hypothalamus during two of the studied age periods (P5-9 and P10-14) (Figure 2). SHR pups had ∼1245 more OX-ir neurons in the hypothalamus than age-matched WKY rat pups at P15, of which ∼74% (∼941) were contributed by newly proliferated BrdU/OX-ir neurons from combined P5-9 and P10-14 neurogenesis, and the remaining 26% were likely contributed by the maturation of existing OX neurons during the same period. It is also important to emphasize that these newly proliferated OX neurons are functional as evidenced by their production of orexin neuropeptide identified via positive immunohistochemical staining for orexin-A, similar to a previous report in 8 week old rats (Xu *et al.*, 2005).

Emerging evidence shows that postnatal and adult neurogenesis are present in the hypothalamus (Xu *et al.*, 2005; Kokoeva *et al.*, 2005; Rojczyk-Gołębiewska *et al.*, 2014; Sousa-Ferreira *et al.*, 2014) and α-tanycytes are likely the neural progenitor cells for generating peptidergic neurons, e.g., orexin neurons, in the hypothalamus (Xu *et al.*, 2005; Lee, 2012; Rizzoti & Lovell-Badge, 2017). Using BrdU, Amiot *et al.* demonstrated that most orexin neurons are generated between embryonic days 11 and 14 (Amiot *et al.*, 2005), while others showed that orexin neurogenesis in the hypothalamus persists post-E14 and into adulthood (Xu *et al.*, 2005;Chang *et al.*, 2012, 2013). Xu *et al.* further showed that hypothalamic BrdU-identified neurogenesis persists into postnatal 8 weeks of age, and they determined that some of the BrdU-positive neurons expressed orexin (Xu *et al.*, 2005).

It is currently unknown what may cause this surge; however, we can speculate that genetic predisposition combined with early postnatal triggers may play a role. Many studies have identified single nucleotide polymorphisms that, when taken together, significantly contribute to hypertension (Pravenec & Kurtz, 2010; Natekar *et al.*, 2014). McCarty and Lee further showed that SHR pups that were cross fostered by WKY dams had significantly lower ABP than SHRs reared by their own mother. On the other hand, WKY pups that were cross-fostered by an SHR dam had no change in their ABP (McCarty & Lee, 1996). These data suggest that while the genetic factors are essential to the development of higher ABP in postnatal development, some aspect of the postnatal development of SHRs is more stressed and serves as a secondary trigger that is also required for the development of a higher ABP. SHR dams have been shown to have increased stress levels, increased tactile stimulation of the pups, altered nutrient exchange during lactation, excessive grooming, and restlessness – all of which could be or contribute to this secondary trigger (McCarty & Kopin, 1978; Tucker & Johnson, 1981; Myers *et al.*, 1989; McCarty *et al.*, 1992; Krukoff *et al.*, 1999). Additionally, chronic stress, e.g. foot-shock, can induce hypertension and double the number of OX neurons in the hypothalamus in rats (Xiao *et al.*, 2013). It is possible that a polygenic predisposition accompanied by prenatal and/or postnatal exposures to increased levels of stress produce an additive effect that leads to the exaggerated OX neurogenesis. The orexin system is associated with hyperarousal, anxiety/stress, and autonomic functions (Kayaba *et al.*, 2003; Furlong *et al.*, 2009; Huang *et al.*, 2010; Johnson *et al.*, 2010; Nattie & Li, 2010) and this early surge of OX activity in SHRs may produce aberrant excitatory drive to many cardiovascular-related nuclei in the brainstem and spinal cord and facilitate the pathological development of hypertension in SHRs.

### Relationship between surge in OX activity and ABP during development in SHRs

The chronological relationship between increased OX activity and ABP in SHRs during development was largely unknown. It is known that ABP increases progressively and rapidly from newborn (∼15-25mmHg) through the first three weeks of life (∼80-90mmHg) in normotensive rats (Zicha & Kunes, 1999). Similar to these previous reports, in this study, we have found that SHR pups have comparable mean ABP and systolic blood pressure (SBP) to the background normotensive WKY rat pups during the first two weeks of life (Figure 3) (Friberg *et al.*, 1989; Dickhout & Lee, 1998; Zicha & Kunes, 1999; Nagai *et al.*, 2003; Li *et al.*, 2013). Around 3-4 weeks of age a measurable difference in ABP between SHR and WKY rat pups emerges and the divergence of ABP between the two strains escalates between weeks 4 and 12 (Lais & Brody, 1977; Friberg *et al.*, 1989; Dickhout & Lee, 1998; Zicha & Kunes, 1999; Nagai *et al.*, 2003). Using a telemetric method in conscious animals, we confirmed that by P20, a small but significant difference in mean ABP began to emerge between SHR and WKY rat pups in both wakefulness and sleep (Figure 3). As discussed above, in SHRs, the number of OX neurons surges to significantly higher than normotensive controls by P15-16, which is about five days prior the emergence of the difference in mean ABP between SHR and normotensive WKY pups (Figure 1 & 3). Even though at this point it is unclear the exact mechanism of such a chronological relationship between increased OX activity and ABP, the closely associated sequential events suggest a potential causal role for the OX system in developing hypertension in SHRs. We speculate that the early surge of orexin signaling may provide excess, and necessary, excitatory drive to many cardiorespiratory-related nuclei in the brainstem and sympathetic preganglionic neurons in the spinal cord to increase SNA, which creates a perfect stage for developing hypertension in SHRs during postnatal development.

### Effects of eliminating excess orexin neurons on ABP and CO_2_ chemoreflex in SHRs

In addition to higher ABP and exaggerated CO_2_ chemoreflex, SHRs also have ∼30% more orexin neurons than normotensive control WKY rats at two ages, P30-58 and adult (Li et al., 2016). Here we further showed that excess OX neurons and higher ABP in SHRs emerges in a sequential order during postnatal development at P15 and P20, respectively. If excess OX neurons are necessary for developing and/or maintaining a higher ABP during postnatal development, then elimination of the excess OX neurons in the hypothalamus may be beneficial in preventing or reducing the aberrant increase in ABP during this period. To test this hypothesis, we selectively eliminated some of the OX neurons using Hcrt-SAP in a subset of SHRs from P25-28, and compared their ABP and CO_2_ chemoreflex with non-lesioned SHR controls 12 days post-lesion at P37-40. The result that eliminating ∼30% OX neurons was sufficient to prevent these OX-lesioned SHRs from developing higher ABP and exaggerated CO_2_ chemoreflex during a postnatal developmental period (P37-40; Figure 4-5), in one way, confirms that excess OX neurons may be necessary in developing higher ABP and exaggerated CO_2_ chemoreflex in SHRs during this developmental period.

On average, the lesioned-SHRs without excess OX neurons had an ABP that was ∼14 mmHg lower than non-lesioned SHRs with excess OX neurons (121 vs 107 mmHg in wakefulness during dark period) at P37-40. The change was similar to that found with OXR blockade in SHRs (Li et al., 2016), where OXR blocker significantly lowered ABP from 121 to 103 mmHg, a level similar to that of WKY rats at the same age (99 mmHg). Although we did not directly compare the ABP of lesioned-SHRs with normotensive WKY rats, our results, combined with previously published reports, suggest that the ABP resultant from excess OX neuron elimination is comparable to the expected ABP of WKY rats at the same age.

In terms of CO_2_ chemoreflex, we previously showed that SHRs have elevated ventilatory and ABP responses to hypercapnia at young (P30-58) and adult ages and that treating with OXR blocker can normalize such exaggerated CO_2_ chemoreflex in SHRs in wakefulness and sleep. Here, we found that the lesioned-SHRs without excess OX neurons had a significantly lower ventilatory response to hypercapnia than non-lesioned rats with excess OX neurons only in wakefulness (Figure 5B). We speculate that the vigilance state-dependency found here could be contributed by 1) a proven effect of Almorexant to promote sleep, 2) possible developmental differences between OX neurons and OXRs during postnatal developmental period, and 3) methodological differences in OX system modulation, e.g. elimination of OX neurons vs blockade of OXRs. Eliminating excess OX neurons will result in loss of other neuropeptides that are also produced by OX neurons, e.g., dynorphin (Chou *et al.*, 2001), even though at this point the role of dynorphin on ABP remains unclear. Of course, a long-term study on the effect of eliminating excess OX neurons or orexin peptide on ABP and CO_2_ chemoreflex in the future may further provide therapeutic significance in human hypertension.

## Additional Information

### Author Contributions

All experiments were conducted at Geisel School of Medicine at Dartmouth. S.B. helped design experiments, performed cell counts, staining, BrdU experiments, and saporin experiments including physiology. RD. L.B., A.M., and J.Y. performed OX and BrdU immunostaining and OX cell counts at different ages; A.L.: designed all the experiments, performed physiology experiments for development data and data analysis. S.B. & A.L. wrote, revised and finalized the manuscript. All authors approved the final version of the manuscript and agree to be accountable for all aspects of the work in ensuring that questions related to the accuracy or integrity of any part of the work are appropriately investigated and resolved and that all persons designated as authors qualify for authorship, and all those who qualify for authorship are listed.

## Acknowledgements

The authors would like to thank Dartmouth undergraduate students Julia Stevenson, Jade Yen, Emily Chen, and Armin Tavakkoli, for their assistance.

